# Assessing bird collisions in the United Kingdom: Modelling frequency of bird-strike from road and rail mortality using a Bayesian hierarchical approach

**DOI:** 10.1101/2020.12.04.412361

**Authors:** Silvia Freire, Lee Read, Todd R. Lewis

## Abstract

Roads are an important way to transport people and goods, but they sometimes have negative impacts on wildlife. One of the leading causes of mortality for several species is identified as road strikes, and the most significant remains bird-vehicle collisions. This study aimed to investigate what species of birds are most affected, and what other factors impact in their susceptibility in road collisions, such as age, sex, season, and type of transports. A total of N=5413 records, and 140 bird species were documented by BTO ringers. For analysis four Bayesian Hierarchical Models were used, with random effects results showing that Barn owls were most affected by collisions. Road mortality presents the highest cause of mortality among species when contrasted with rail mortality. Age and sexual bias was detected across all species, however juveniles and males did appear to be prominent in relation to other age classes. Winter and early spring were the months with most reported casualties and 2016 had lower abundance of mortality across the 10-year period. 75% of birds were found within a week, which may indicate some bias interference from scavenging animals, as true figures could be up to 16 times more. This study discusses some mitigation measures found in current research, that could dramatically reduce numbers of birds affected each year by road mortality.

## Introduction

At present there are over 64 million kilometres of roads on the earth, and Great Britain comprises almost 400,000km of asphalt road, which could circulate ten times around the globe (Cooke, Balmford, Johnston, Newton, & Donald, 2020). While roads are important to human society as a means of transport of people and goods, they can have negative impacts on wildlife (Arnold et al., 2019; Johnson, Evans, & Jones, 2017; Meijer, Huijbregts, Schotten, & Schipper, 2018).

Wildlife can suffer adverse effects from roads, for example by direct impacts like collision with transport, or by indirect impacts like the fragmentation of habitats (Schwartz, Williams, Chadwick, Thomas, & Perkins, 2018). In addition, roads contribute to negative effects by noise, pollution, and light (Johnson, Evans, & Jones, 2017).

Roads can be ecological snares for certain species, as an appealing habitat or source of food they may encourage animals to them, only for them to subsequently endure health effects, decreased reproductive success or vehicle-collision as consequence (Cooke, Balmford, Johnston, Newton, & Donald, 2020).

Over the last century, the effects traffic has on the survival of animals has been notable (Møller & Erritzøe, 2017). One of the leading causes of mortality for several species is identified as collisions amongst vehicles and wildlife (Gonzalez-Suarez, Zanchetta Ferreira, & Grilo, 2018). The most considerable remains bird-vehicle collisions, with estimations in some countries of 80-340 million in the United States, 57 million in Europe, and 27 million in England (Husby, 2017; Loss, Will, & Marra, 2015; Møller & Erritzøe, 2017).

Regardless of the considerable amount of investigation into such numbers, insufficient research has investigated the effects of roads on birds, possibly because of misconceptions of flight capabilities that allow them to escape and thereby avoid related effects (Johnson, Evans, & Jones, 2017). Species of birds more profuse alongside roads are usually more often implicated in accidents with vehicles (Madden & Perkins, 2017). Variations in mortality trends might be propelled by alterations in roadkill coverage techniques or driving performance (Madden & Perkins, 2017). Mortality likelihood might also be partially a result of birds’ cognitive skills (Møller & Erritzøe, 2017). There is a need to take into consideration all such factors by assessing changes in mortality rate of different bird species during extended periods (Madden & Perkins, 2017).

### Avian Senses and Behaviour

Birds constitute an integral part of a complex network in the environment, preserving and supplying several ecological provisions which humans are reliant on for continuous development and success (Johnson, Evans, & Jones, 2017). Birds are ecological facilitators, as they pollinate and disperse important plant species, and they also predate on species that are considered pests across agricultural industry (Johnson, Evans, & Jones, 2017).

The susceptibility of specific species to road traffic is dependent on their behaviour and environment (De Jong, van den Burg, & Liosi, 2018). The information that birds obtain visually from their environment is clearly distinct from that obtained by humans under similar conditions (Martin, 2011). The reason for this is the basic dissimilarities among birds and primates on all degrees of structure of their optical systems, involving physiological optics, retina, visual information they obtain by the brain, and visual areas (Martin, 2011).

Birds’ senses display a high level of deviation which seems to be responsive to the cognitive challenges presented, particularly hunting, and thus sensory abilities are perceived as an essential part of every species’ ecosystem (Mitkus, Potier, Martin, Duriez, & Kelber, 2018). Birds use vision as their main sensory organism, with many species sharing ultraviolet spectrum (May, Åström, Hamre, & Dahl, 2017). Bird colour vision is resolved by photoreceptor types, therefore some bird’s behaviour is driven by UV indications (Lind, Mitkus, Olsson, & Kelber, 2013).

Owls and diurnal birds of prey have bigger eyes when compared to other birds that fly (Mitkus, Potier, Martin, Duriez, & Kelber, 2018). The visual sharpness in a few diurnal birds of prey is a result of their larger eyes, although in contrast to nearly all other species, they are not especially responsive to ultraviolet light (Mitkus, Potier, Martin, Duriez, & Kelber, 2018). Statistical evaluation of every existing visual field information has demonstrated that the binocular fields of non-passerine birds are considerably smaller than passerine species (Mitkus, Potier, Martin, Duriez, & Kelber, 2018).

Dangers to birds of prey, and the way they react visually, mostly derive from predator avoidance, but also strongly from human interaction. They are predisposed to evading predation and collision, for which transportation collisions is the most frequent (Hawk Watch International, 2018). Birds of prey often forage on roads, searching for prey on the edges of roads, even if they effectively catch their prey, they can collide with transportation when attempting to fly off while carrying heavier animals (Hawk Watch International, 2018). When birds of prey are foraging, they fly down with extreme focus, described as tunnel-vision, as they predate on prey, at times flying across roads when they can get hit by vehicles (Hawk Watch International, 2018). Owls are extremely susceptible to road mortality due to their lower hunting flight heights (De Jong, van den Burg, & Liosi, 2018).

### Road Design and Infrastructures

Urban development influences migration patterns and wildlife dispersals (Morelli, Beim, Jerzak, Jones, & Tryjanowski, 2014). There is considerable research showing the environmental effects of roads, which turn out to be a major threat to species richness, supporting a significant reduction in bird populations in the vicinity of roads, in particular where traffic is abundant (Selva et al., 2011; Cooke, Balmford, Johnston, Newton, & Donald, 2020). Birds are particularly useful in assessing the effects of anthropogenic disturbance, with road and railway networks already well known as causing adverse effects (for example; sound levels, habitat destruction, obstacle impacts, disturbance and mortality from accidents (Morelli, Beim, Jerzak, Jones, & Tryjanowski, 2014)).

In Great Britain, several bird populations have had a significant reduction over past years, these decreases were associated to many influences, such as: global warming; shifts in land use and farming practices, habitat destruction and fragmentation (Cooke, Balmford, Johnston, Newton, & Donald, 2020). Additionally, roads might have added to these population declines, as traffic quantity has risen by more than 160% in the last sixty years (Cooke, Balmford, Johnston, Newton, & Donald, 2020; Roulin, 2020).

Since 1960, there has been a rise in speed and traffic quantity, and road collision levels are dependent on these two factors (Madden & Perkins, 2017; Orlowski, 2008). Subsequently, according to Orlowski (2008), the English population of House sparrow *Passer domesticus* has decreased by 13% due to road mortality. Other recent studies suggest road accidents to be a primary cause of mortality amongst the decreasing populations of owls in the countryside, such as Barn owl *Tyto alba* and Little owl *Athene noctua* (Orlowski, 2008).

Millions of birds are dying every year from traffic collisions, however, highway schemes usually concentrate on decreasing road collisions with larger mammals due to security and financial motives (Arnold et al., 2019). Presently, ever-increasing new hedgerows are being planted as an integral part of conservation and agricultural programs, which set ideal environments for some species in decline (Orlowski, 2008).

Numerous suggestions on prevention measures of bird road mortality are discussed in various papers; open verges and grassland located close to roads being recommended to grow into shrubs, in order to reduce accessibility to foraged prey such as rodents (Figure 1A) (Barn Owl Trust, 2015; Orlowski, 2008).

**Figure 1.**
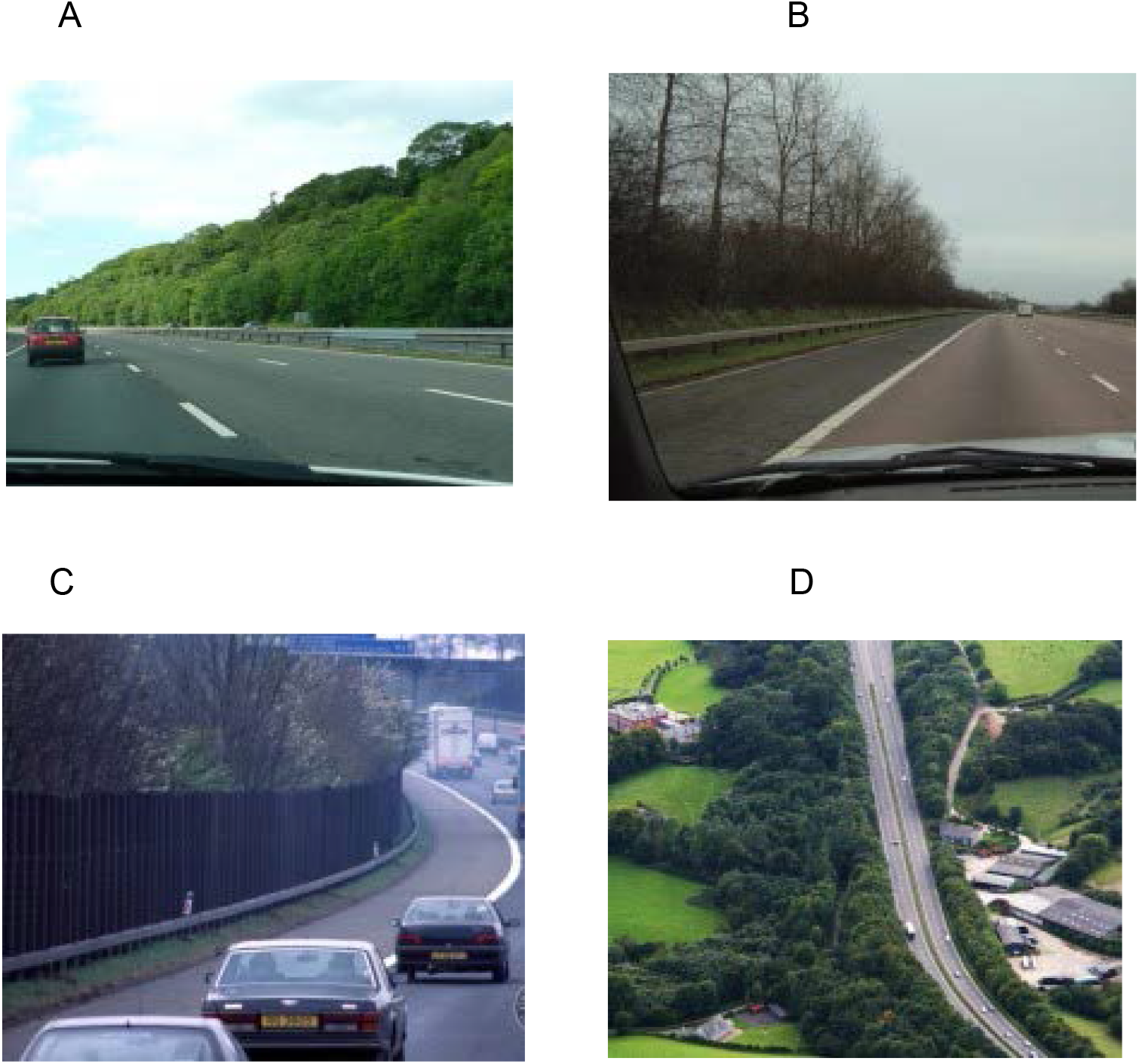
Some mitigation options currently suggested for birds of prey include shielding main roads: (A) edge scrub is an inexpensive alternative, directing birds to fly higher (between 2-5 metres should help to avoid lorries) as well as supporting wildlife and allowing drivers to slow down in the event of colliding; other more costly alternatives consist of (B) tall trees and (C) wall barriers; and (D) a secure area with trees on either side (Barn Owl Trust, 2015).

Another method can be to make birds fly higher, by placing fences and lines of tall trees closer together (above 3 m) in addition to building walls (Figure 1B and 1C) (Orlowski, 2008). Planning roads with such barriers may help to divert owls and birds of prey as they will fly higher, and therefore could reduce collisions between certain species (Figure 1D) (Ramsden, 2003; Boal & Dykstra, 2018; Roulin, 2020).

Additional mitigation measures include decreasing the quantity and speed of transportation and raising the awareness of drivers. This has been demonstrated to enhance animal numbers in regions with abundant road system, although attempts to employ these shifts have been frequently challenging (Boal & Dykstra, 2018; Selva et al., 2011).

In summary, there is mounting evidence that road collisions are a major threat to bird populations. This study aimed to evaluate what species are most affected by road mortality and what transportation type has a greater impact. To understand what factors might influence road collisions trends factors such as bird age, bird sex, month, or year of mortality, number and species affected, and conditions the birds were found in were studied. This work sought to investigate relationships between age, sex, month, year, and types of transportation in relation to the abundance (and frequency) of bird collisions.

## Methodology

Data used in this study was provided by the British Trust for Ornithology (BTO), comprising records of reported birds in road-collision. The data collected was supplied by ringers throughout Great Britain and Ireland, reporting ringed birds, which may have either a metal ring, metal and colour ring, or less frequently just a colour ring. The BTO Ringing Scheme is funded by a partnership of the British Trust for Ornithology, the Joint Nature Conservation Committee (on behalf of: Natural England, Natural Resources Wales and Scottish Natural Heritage and the Department of the Environment Northern Ireland), The National Parks and Wildlife Service (Ireland) and the ringers themselves.

The data comprised N=5413 records of 140 species involved in road collisions over the past ten years. It also included the types of transports implicated, and other relevant information such as age, sex, month, year, and encounter conditions. Statistical models were used to uncover relationships between species, factors, and collision trends. The factors codes (Age and Finding Condition) are described in – Appendix 1.

### Data Analysis

For the statistical analysis four Bayesian hierarchical models with random effects were structured into subsets of broad-based bird guilds; Raptors (Subset 1), Seabirds (Subset 2), Wildfowl (Subset 3), and Garden Birds (Subset 4). Data was converted first using package dplyr (Wickham et al., 2020) piping categorical replicated data rows into numerical integer counts as response variables. Estimation was conducted using Markov Chain Monte Carlo (MCMC) routines in software JAGS version 4.3.0 (Plummer, 2003) through the package runjags (Denwood, 2016) in R version 3.6.3. (R Core Team, 2019).

Models were expressed similarly to a frequentist log-linear model used in lme4 as;

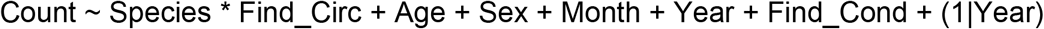

Each model was fitted as;

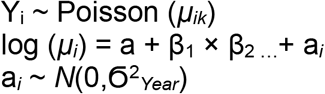

where; k is the over-dispersion parameter, a is intercept, β_1_ and further β are fixed effect coefficients, and a_*i*_ is the randomized effect (Year).

Models comprised Poisson family distributions running 40000 iterations, 10000 discarded for burn-in, and 4 chains. Priors were set using templates within runjags modest automated uniform gamma distribution detected and set through JAGS (priors = ∼ dnorm (0, 10^-6)). Convergence was assessed using MCMC trace plots of iterations retrieved from runjags and inspection of the Gelman-Rubin statistic potential scale reduction factor (psrf) (Gelman et al., 2013).

Model assumptions of mean-variance, log-linearity and potential autocorrelation were explored using residuals vs. fit plots, and a correlation plot function within runjags. MCMC draws from posterior distributions were used for evaluating model component relations. Factor interactions plots were constructed in ggplot2 (Wickham, 2016) to contrast model findings. Summary data was recovered from base-R functions.

## Results

Summary results across all data are shown in Table 1. Convergence, and autocorrelation are reported in – Appendix 2. Convergence was complete for all variables in all four subset models 1-4. Autocorrelation levels were acceptable with various levels of only minor correlation across some factors. Models of Subsets 1-4 generated a series of results, much of which demonstrated interesting and diverse variation in response to encountered circumstance (found circumstance) Find_Circ (Road or Rail related mortality) (Appendix 2).

**Table 1.** Summary of N bird strike across model subsets: dominant species are shown

| SUMMARY DATA |  |  |  |  |  |  |  |  |
| --- | --- | --- | --- | --- | --- | --- | --- | --- |
| Factor | Model Subset 1 |  | Model Subset 2 |  | Model Subset 3 |  | Model Subset 4 |  |
| Species | BarnOwl | 2134 | HerringGull | 138 | MuteSwan | 171 | Blackbird | 522 |
| Species | Kestrel | 191 | Oystercatcher | 109 | Mallard | 53 | HouseSparrow | 113 |
| Species | TawnyOwl | 187 | LesserBlack-backedGull | 65 | CanadaGoose | 20 | Greenfinch | 104 |
| Species | Buzzard | 94 | Black-headedGull | 40 | Lapwing | 20 | Bluetit | 103 |
| Species | Sparrowhawk | 68 | CommonGull | 16 | Coot | 16 | Chaffinch | 96 |
| Species | RedKite | 61 | GreatBlack-backedGull | 14 | Curlew | 15 | Goldfinch | 84 |
| Species | (Other) | 109 | (Other) | 29 | (Other) | 81 | (Other) | 760 |
| Age | UNK | 2422 | UNK | 334 | DHY | 15 | UNK | 1099 |
| Age | JUV | 177 | DHY | 24 | FGH | 3 | JUV | 318 |
| Age | DHY | 91 | SEC | 14 | JUV | 17 | SEC | 222 |
| Age | SEC | 71 | THR | 14 | SEC | 33 | DHY | 72 |
| Age | FGH | 67 | JUV | 13 | THR | 8 | FGH | 66 |
| Age | THR | 13 | NSTUNK | 5 | UNK | 300 | NST | 3 |
| Age | (Other) | 3 | (Other) | 7 | NA | NA | (Other) | 2 |
| Sex | F | 196 | F | 1 | F | 33 | F | 204 |
| Sex | M | 219 | M | 5 | M | 46 | M | 480 |
| Sex | U | 2429 | U | 405 | UNK | 297 | UNK | 1098 |
| Month | JAN | 551 | JUL | 103 | 0 | 0 | MAY | 320 |
| Month | MAR | 378 | JUN | 72 | 0 | 0 | JUN | 272 |
| Month | FEB | 358 | 0 | 0 | 0 | 0 | APR | 267 |
| Month | APR | 244 | 0 | 0 | 0 | 0 | JUL | 186 |
| Month | AUG | 224 | 0 | 0 | 0 | 0 | MAR | 178 |
| Month | SEP | 224 | 0 | 0 | 0 | 0 | JAN | 150 |
| Month | (Other) | 865 | (Other) | 71 | (Other) | 124 | (Other) | 409 |
| Year | 2015 | 358 | 2013 | 60 | 2010 | 53 | 2010 | 221 |
| Year | 2010 | 318 | 2018 | 49 | 2011 | 44 | 2011 | 219 |
| Year | 2011 | 315 | 2017 | 43 | 2015 | 42 | 2012 | 203 |
| Year | 2012 | 309 | 2014 | 42 | 2016 | 41 | 2013 | 187 |
| Year | 2018 | 309 | 2012 | 40 | 2014 | 37 | 2017 | 180 |
| Year | 2013 | 308 | 2010 | 39 | 2012 | 36 | 2014 | 172 |
| Year | (Other) | 927 | (Other) | 138 | (Other) | 123 | (Other) | 600 |
| Find_Cond | AWU | 106 | AWU | 21 | AWU | 11 | FDW | 1480 |
| Find_Cond | DNF | 191 | DNF | 19 | DNF | 20 | DNI | 143 |
| Find_Cond | DNI | 246 | DNI | 29 | DNI | 23 | DNF | 73 |
| Find_Cond | DYG | 111 | DYG | 14 | DYG | 13 | DYG | 44 |
| Find_Cond | FDW | 2020 | FDW | 254 | FDW | 275 | AWU | 15 |
| Find_Cond | FDWDNI | 126 | FDWDNI | 67 | FDWDNI | 30 | FDWDNI | 12 |
| Find_Cond | SWU | 44 | SWU | 7 | SWU | 4 | (Other) | 15 |
| Find_Circ | Rail | 123 | Rail | 5 | Rail | 25 | Rail | 8 |
| Find_Circ | Road | 2721 | Road | 406 | Road | 351 | Road | 1774 |

A significant amount of count response species across models did not exhibit a lot of variation in differences (Appendix 2). This is mainly caused by the sparse data across the models or limited sample size (for instance, Subset 2 - Arctic tern *Sterna paradisaea*, N=3). Other species responses were well represented (for example, Subset 1 – Barn owl *Tyto alba*, N=2134; Table 1). Caterpillar bar plots with confidence intervals are shown for all species (Figures 2-4). Species that had interaction and were strongly affected as a result or Road or Rail mortality are those with clear positive or negative difference from the mean zero line and particularly those with confidence intervals that do not have zero crossings.

**Figure 2.**
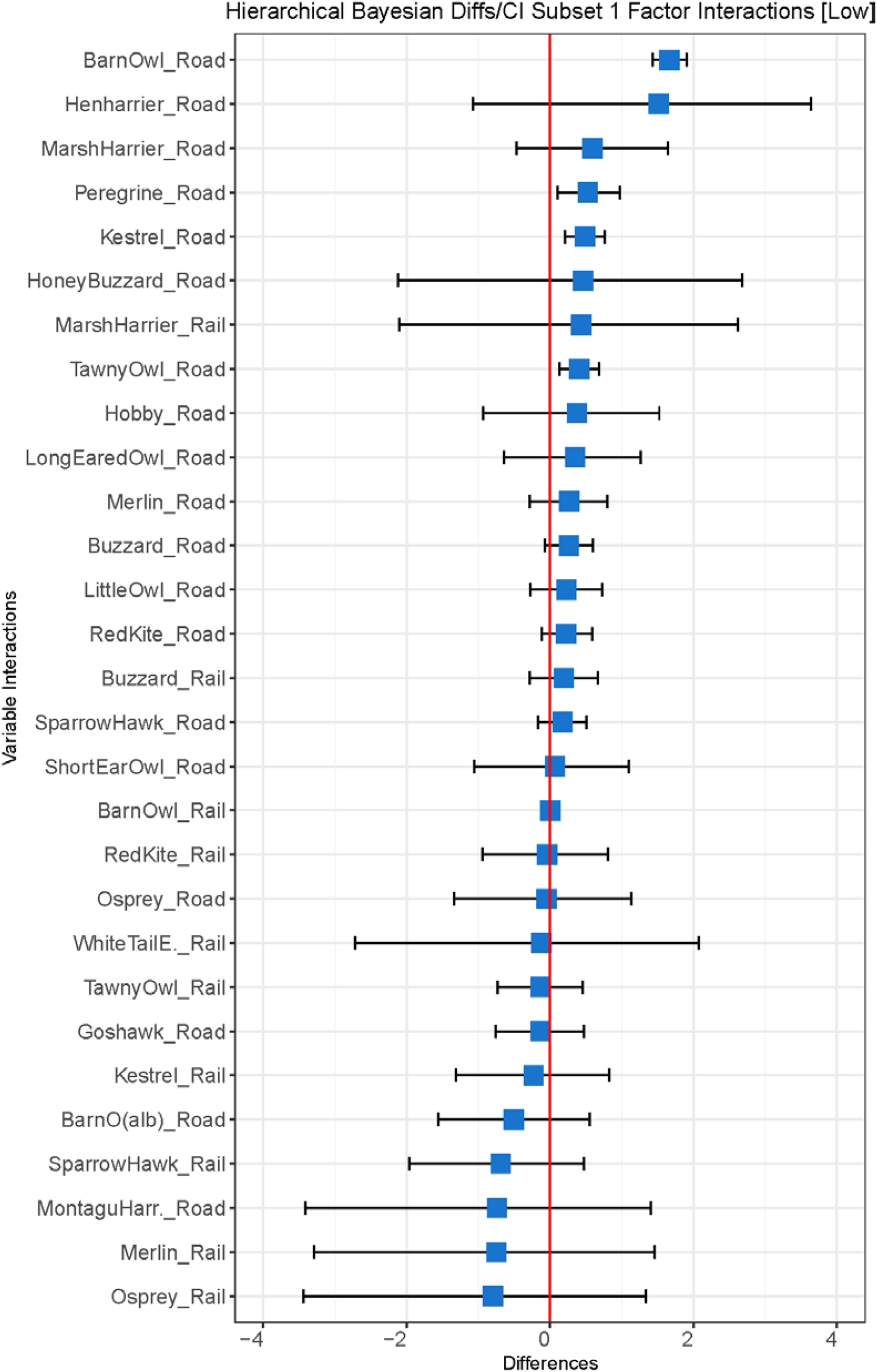
Caterpillar bar plots with credible intervals for Subset 1 species (Note, Barn Owl, Kestrel, Peregrine, Tawny Owl and Buzzard on Roads). Species with CI clear of the zero crossing differences are highly impacted

**Figure 3.**
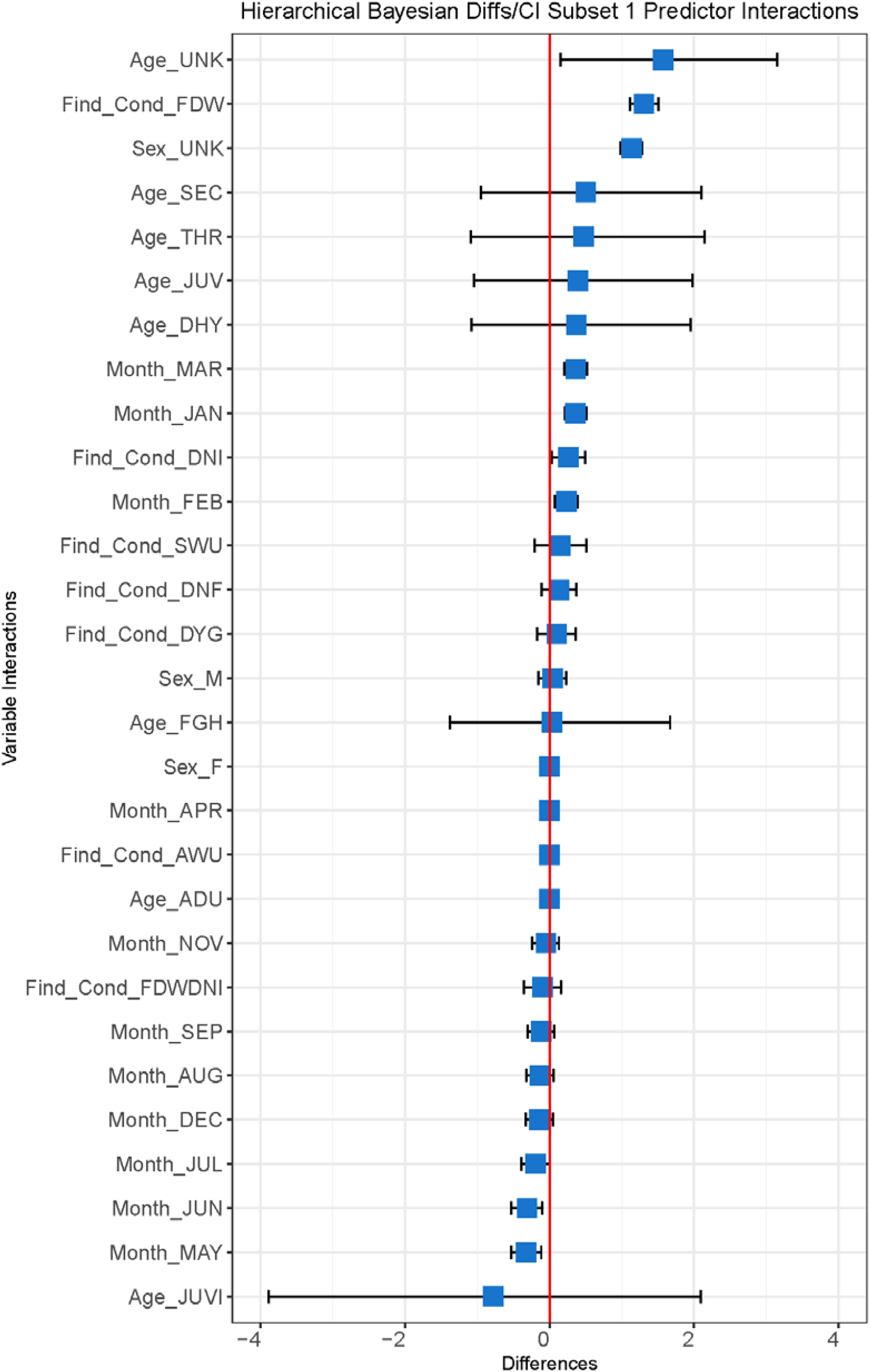
Caterpillar bar plots with credible intervals for Subset 1 predictor levels (Note, Find_Cond_FDW/DNI, Sex UNK, and January, February, March, May and June). Factors with CI clear of the zero crossing differences are influential

**Figure 4.**
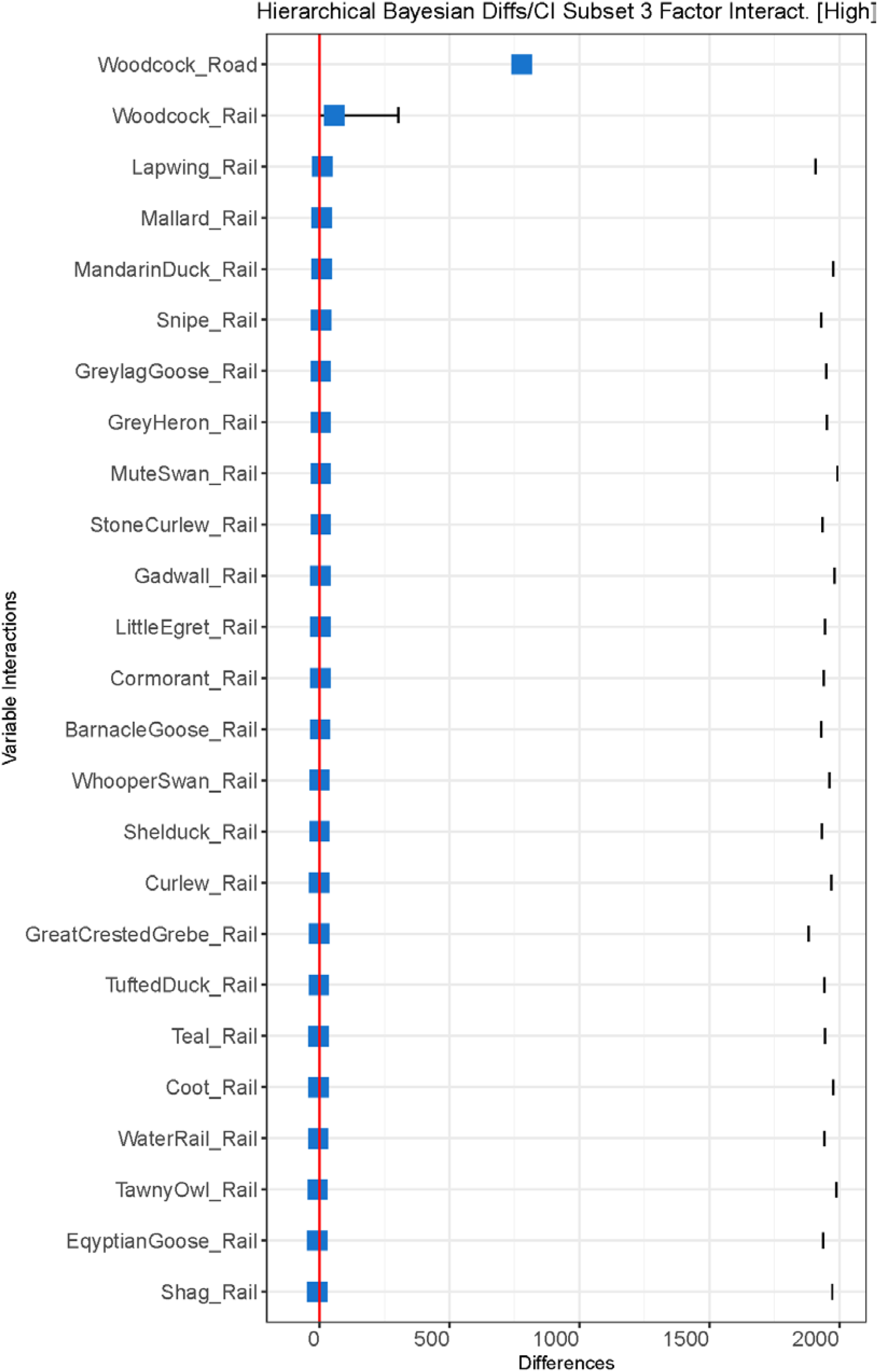
Caterpillar bar plots with credible intervals for Subset 3 species (Note, Woodcock in both Rail and Road). Species with CI clear of the zero crossing are influential. Detached CI’s with mean values (blue) on the zero line are non-significant

Few structured hierarchical level predictors (Age, Year, Find_Cond…) exhibited strong interactions. However, in Subset 1 – Raptors were greatly affected and demonstrated an obvious impact of a seasonal effect on mortality frequency for some species (JAN, FEB, MAR, MAY), sexual bias (UNK), and age related influence (UNK). For Subset 1, Barn owls, Kestrels *Falco tinnunculus*, Peregrines *Falco peregrinus*, and Tawny owls *Strix aluco* were clearly affected worst by Roads, a concerning trend (Figure 2). Woodcock *Scolopax rusticola* in Subset 3 were impacted for both Rail and Road mortality (Figure 4). Buzzards *Buteo buteo* had expressed confidence intervals on/just over the zero crossing and presumably with further data would represent a significant impact from Road collisions. Overall, there are clear seasonal effects on one guild (raptors) with casualties on Roads, and a lack of age and sexual identification from specimens of some species (Figure 3).

In all subsets, the species most affected by road collisions (by both Road and Train) were Barn owls and Blackbirds *Turdus merula* (Figure 5). Age bias was shown for all the other species (UNK – 77%) presented as the highest followed by (JUV – 10%) across the age range (Figure 6). Sexual bias was also shown across all species (UNK – 78%), (M – 14%) in comparison with (F – 8%) (Figure 7). In every subset, the month with highest casualties was (JAN) followed by (MAR and APR) (Figure 8), and across years illustrating more range from 2010-2013, while 2016 was the year with lower casualties reported (Figure 9).

**Figure 5.**
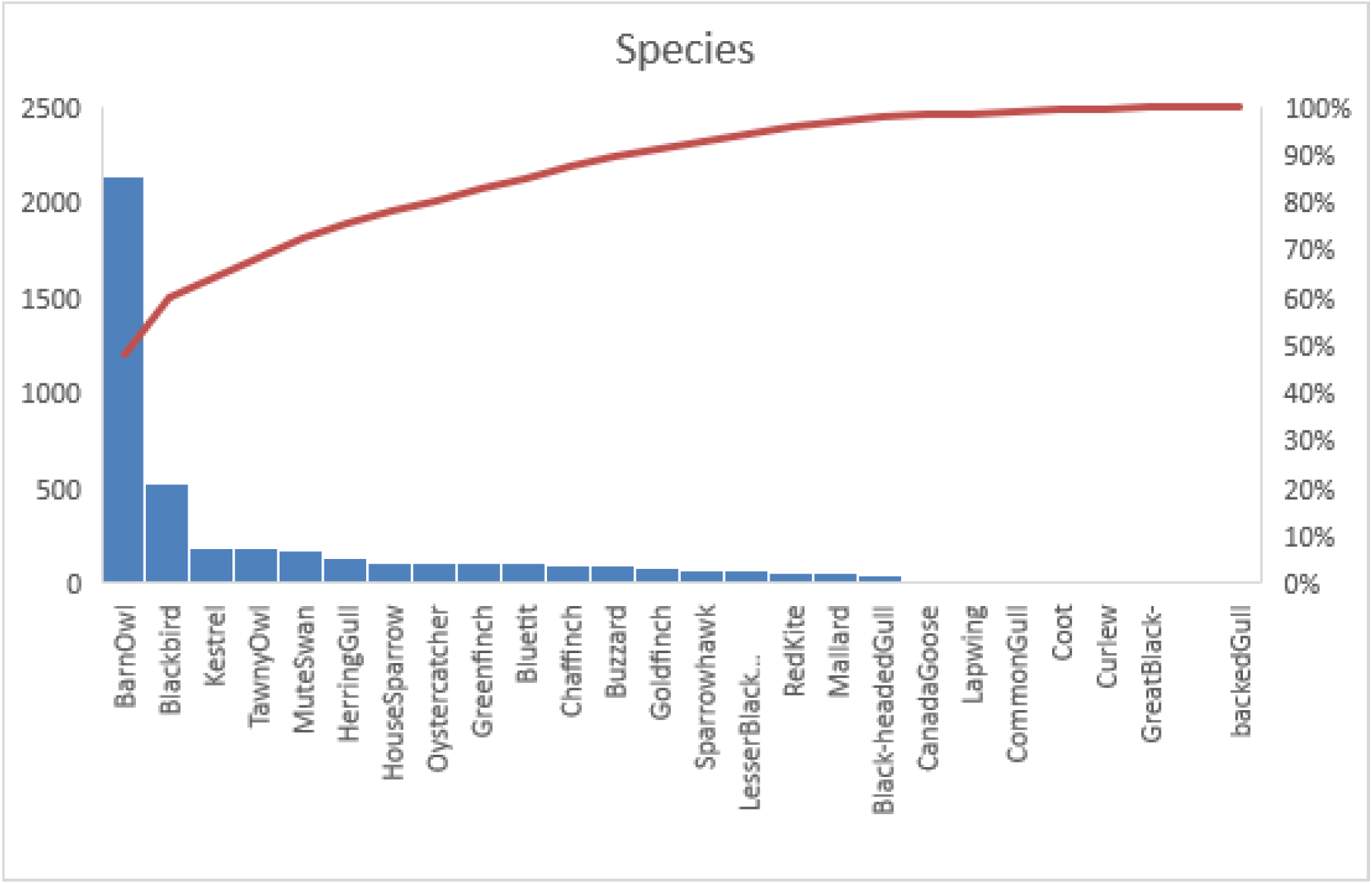
Summary chart showing species highly impacted by road/rail collision - all subsets

**Figure 6.**
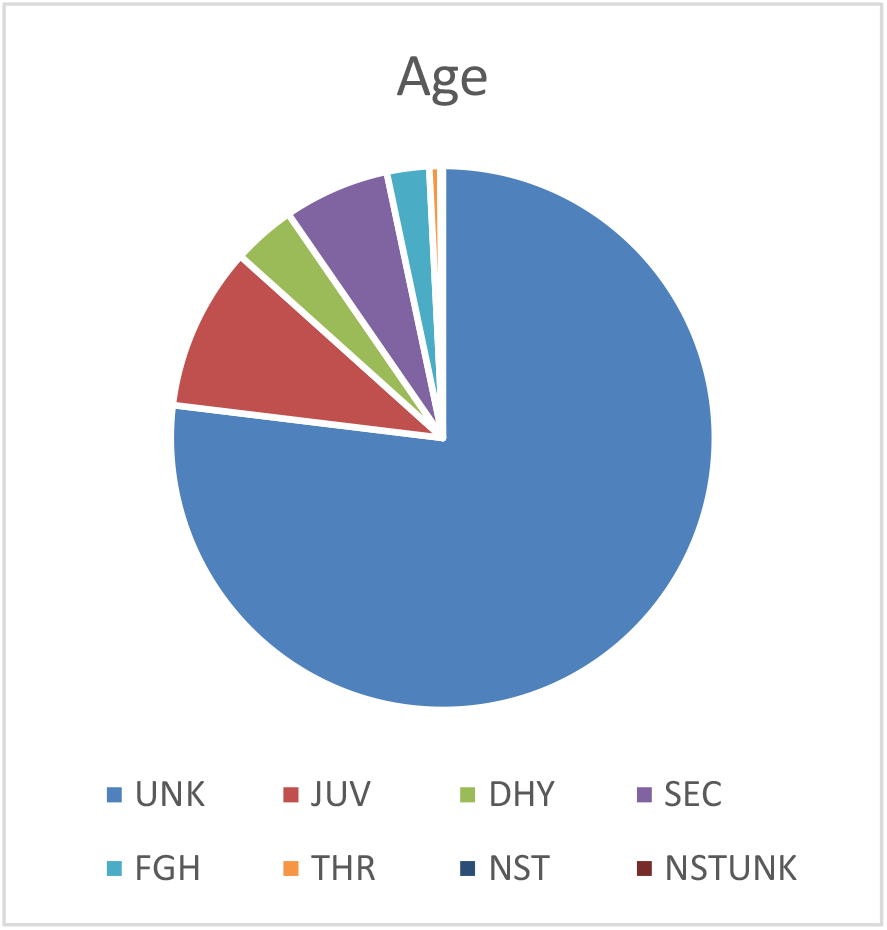
Pie chart describing age range

**Figure 7.**
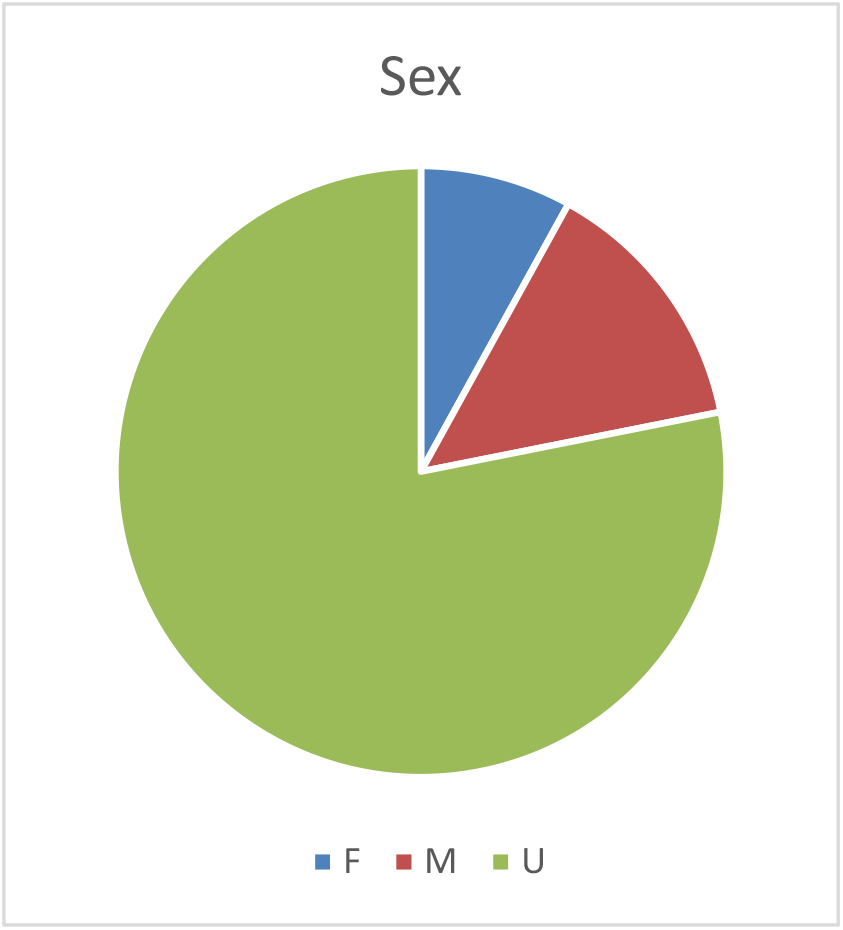
Pie chart showing gender

**Figure 8.**
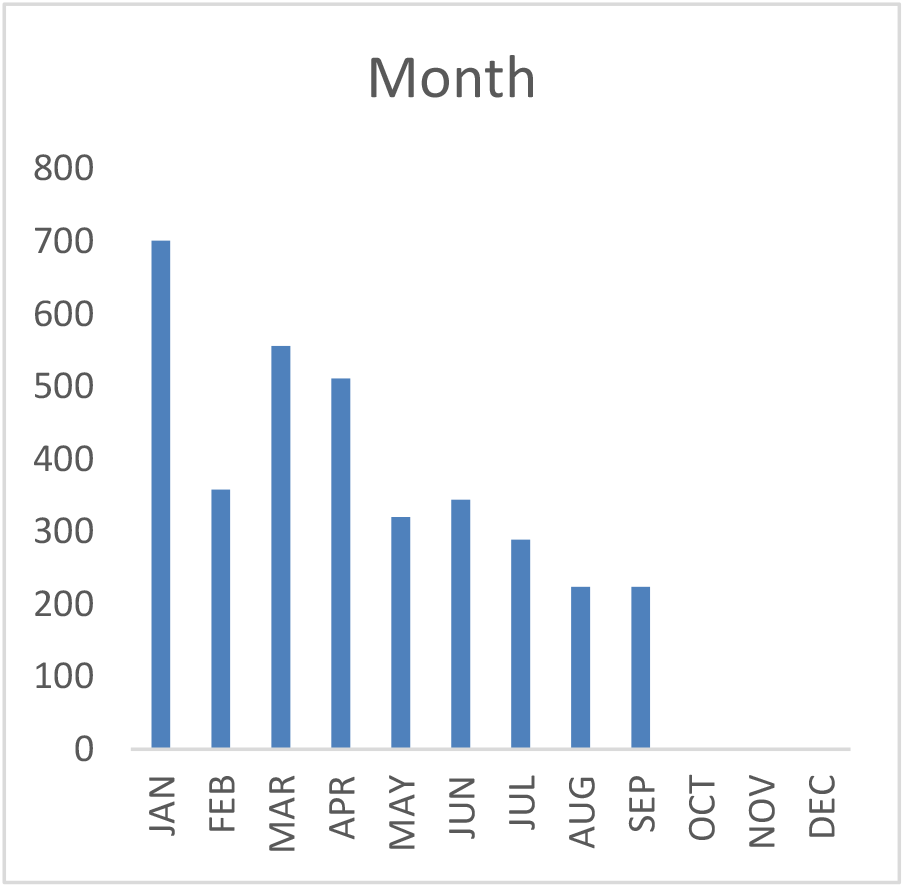
Summary of mortality by months

**Figure 9.**
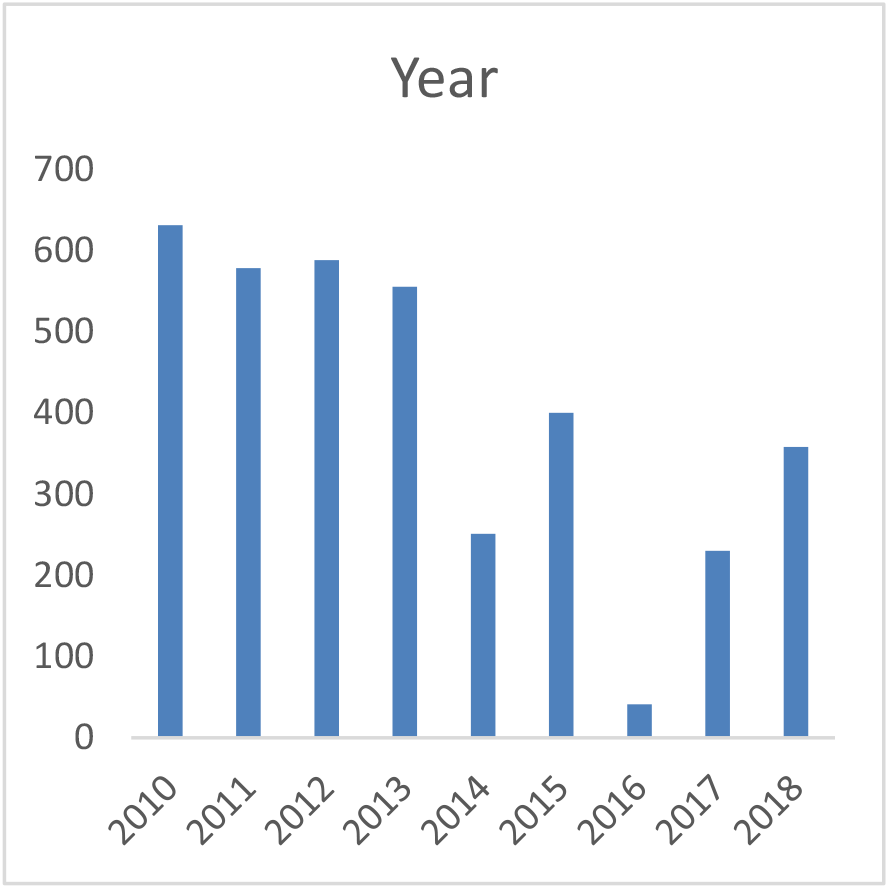
Summary of mortality by years

The Finding Conditions of all species suggested a higher trend for (FDW – 75%) (Figure 10). Rail related mortality was expressed less across all Subsets and species (Figure 11) and perhaps may benefit from further, separate, investigative modelling for more expression.

**Figure 10.**
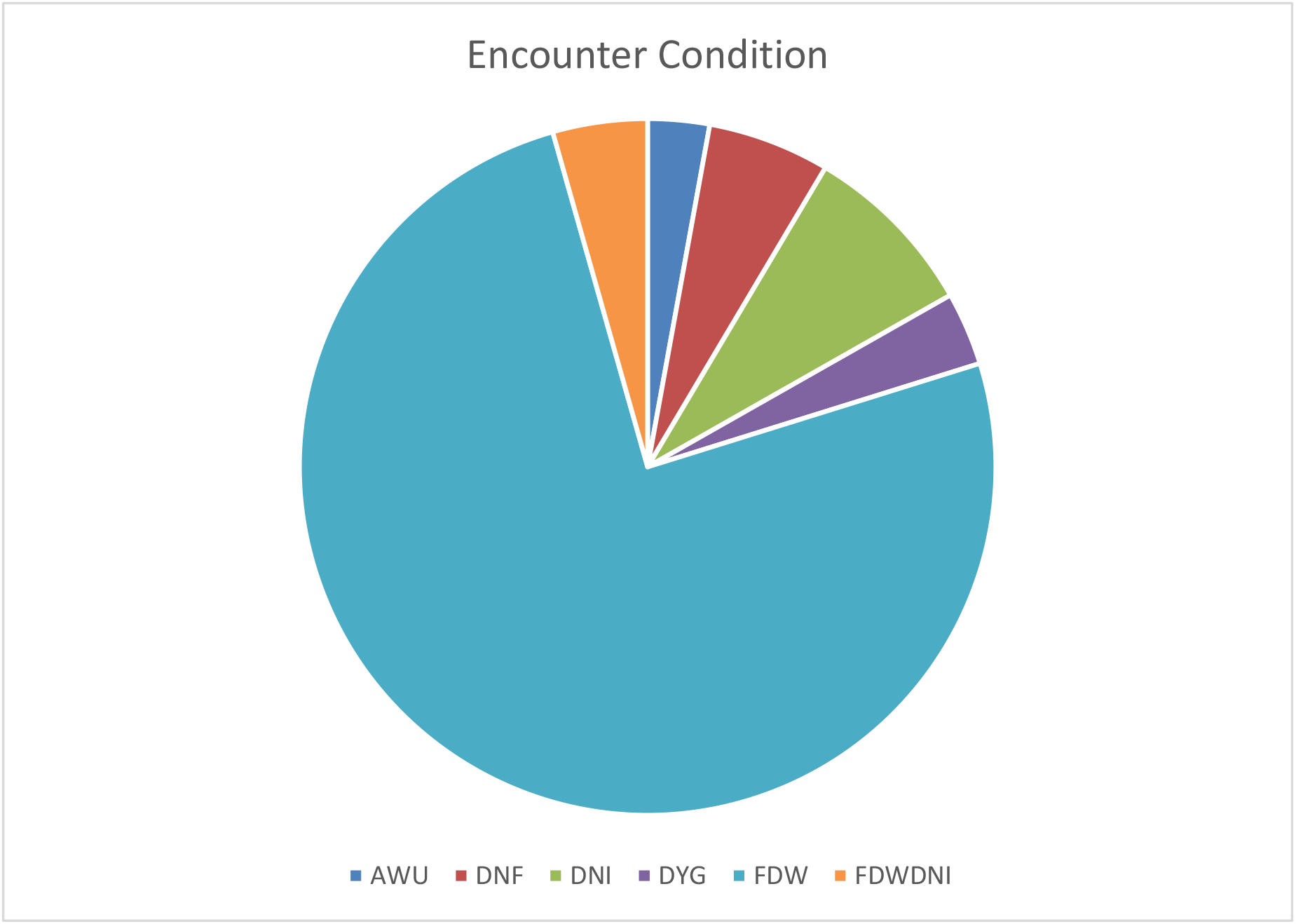
Pie chart showing encounter conditions found

**Figure 11.**
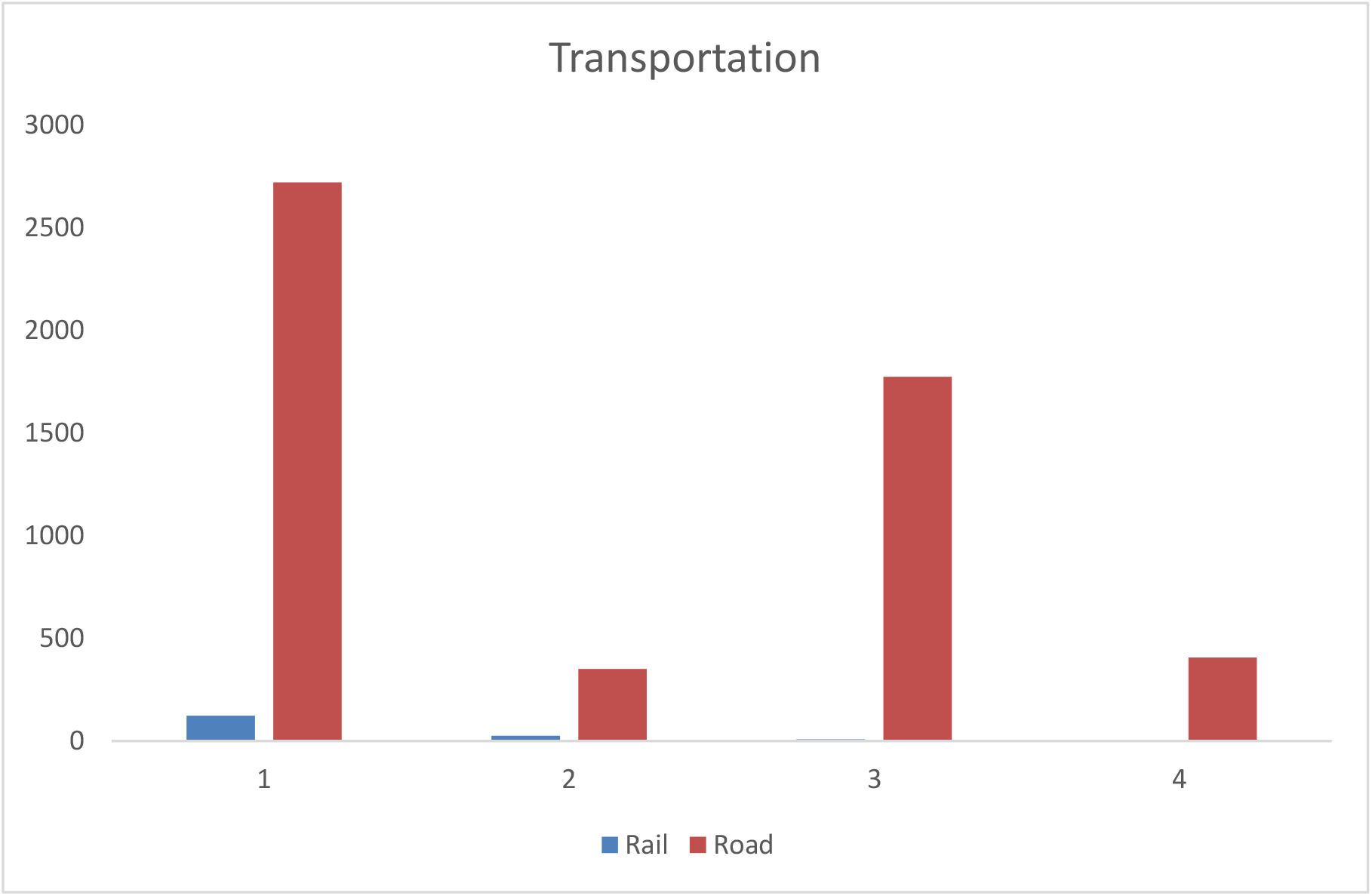
Summary chart of types of transportation influencing bird collision

## Discussion

The probability of collisions with transports varies between the groups of birds (Husby, 2017). Across all Subsets (Raptors, Seabirds, Wildfowl and Garden Birds), Raptors were the most impacted by vehicle collisions, followed by Garden Birds. Raptors are very reliant on road environments as they provide them with food (Husby, 2017; Kajzer-Bonk et al., 2019). Barn owls were particularly affected within results of this study, supporting other studies that reached similar conclusions. Barn owls are often implicated in vehicle collisions, which constitutes a persistent pattern and may have an impact on populations of this species (Arnold et al., 2019; Cooke, Balmford, Johnston, Newton, & Donald, 2020). Arnold et al. (2019) has shown that Barn owl transportation collisions are less expressed on roads with shrubs when contrasted with roads with grass on their edges; thus collisions are believed to be a result of the concentrations of small mammals near the edges of roads in various areas.

In this study, it was not statistically possible to make inferences between different sites to understand areas of foraging preference, as spatial data was not provided, however, this could be a potential factor influencing the frequency of Barn owl casualties.

Blackbirds also showed high numbers impacted by transport collisions. They are perhaps not as adversely affected as Barn owls, but are still higher in frequency than all the other species across the Subsets. Roadside surroundings are appealing for this species, for breeding and foraging, and they often perch on trees and use shrubs for hunting nearby roads (Husby, 2017).

The age of almost all the birds reported were biased with more unknown (UNK) reports than known aged assessment. This is understandable given the difficulty of age class assessment for birds killed by impact. Notwithstanding, some UNK reports are followed by JUV, which shows that young birds (juveniles) are greatly impacted when compared to with other age ranges. Road collisions continue to be a significant cause of mortality among young birds, especially after breeding season, as they fledge their nest and disperse away from the place they were born to create their own territory (British Trust of Ornithology, n.d.). Young birds, which have less spatial experience, might be drawn to roadsides by the potential for food resources, which expose them further to transport collisions (Kajzer-Bonk et al., 2019).

Sex ratio was biased (UNK) across all the species. However, males (M) presented more frequency that female birds (F), which might be an artefact within the data as males were more frequently reported. Ecologically, this could be due to males foraging for longer periods of time, especially during the breeding season, whereas females often spend more time in incubation and feeding of offspring (De Jong, van den Burg, & Liosi, 2018).

Seasonal changes may be able to justify the difference in the number of birds reported from roads (Husby, 2017). January presented as the month with highest numbers of casualties, followed by March and April. These results suggest that bird collisions were higher during winter and early spring (breeding season). This could be due to scarcity of prey in winter, and thus the need for birds to forage on roads during this time. Also, higher frequency of reported birds during March and April indicated there was a tendency for more collision during the breeding season. Prey availability could influence trends across years, as there are years with considerably higher numbers. Year 2016 had the lowest casualties. In years that prey was abundant, collisions were greatly decreased (De Jong, van den Burg, & Liosi, 2018).

Mortality was greatest among birds that were found dead within a week, which indicated that birds that are found in a short period of time are more likely to be reported. Studies on bird collisions may be influenced by numerous biases due to prompt scavenging, thus when adjusting for this, mortality rates could be considered up to 16 times higher (Madden & Perkins, 2017; Guinard, Julliard, & Barbraud, 2012). Further, unbiased studies are needed to assess the connection between mortality rates, surrounding environment, and road structure (Guinard, Julliard, & Barbraud, 2012).

Road mortality is clearly dominant in casualties when contrasted with railway transportation, indicating that roads are possibly more appealing to birds than railways. This could be due to road design or abundance of prey. Also, the edges on roads and shrubbery have significant capacity as bird territory (Dover, 2019). Barn owls are drawn to grassy fields because of larger quantities of prey (Arnold et al., 2019; Barn Owl Trust, 2015). To potentially mitigate for this, zones of wild grass could be supplied closer to roads, but only if such areas are screened (Ramsden, 2003).

Meticulous design of roadside flora assemblages is therefore crucial to decrease negative impacts in order for birds and wildlife to benefit (Dover, 2019). To decrease owl road mortality, perhaps a realistic choice of mitigation would be to further implement strategies and management to enhance the quality of farmlands in such a manner that alternative land would be able to support larger numbers of prey (rodents) (Arnold et al., 2019; De Jong, van den Burg, & Liosi, 2018).

The most appropriate practice of bird protection around roads may well be to prevent unintentional invitation of birds toward them (Arnold et al., 2019). Decreasing or removing vegetation, often removing carcasses, so that such areas are not so attractive to foraging birds, or applying road lightning with colours and patterns to try to dissuade birds from being drawn to them (Arnold et al., 2019) are all useful strategies. Ultimately, further investigation is required to understand relationships between location, number and species affected by road collisions.

## Conclusion

This study has shown that road mortality among birds is considerably high, across a variety of species, particularly Barn owls. This could be due to their behaviour and foraging opportunities. Also, in winter and early spring mortality frequency is higher possibly due to the lack of prey availability which could encourage birds to forage around roads.

Also, during breeding season, as juvenile birds explore their territory range, and because they are spatially inexperienced, they may forage more frequently on roads. Collisions are more frequent on roads than railways, which suggests that roads pose an attractive environment for birds. This could also be a result reflected in a greate frequency of reporting along roadside environments. Considering most dead birds were found (detected) within a week, scavenging may be taking place naturally, that may not allow for accurate abundance of road or rail mortality reporting.

Road surroundings should be carefully planned and designed, providing increased foraging areas, especially for Barn owls, which are greatly affected by road mortality. This, if applied correctly, could help Barn owls to divert from using these areas. The screening of roads (shrubs, trees, or panels) could help to decrease such road mortality, as birds would have to fly higher and this would subsequently prevent increased bird collisions. Eliminating or modifying grassy road edge areas could decrease accessibility by small mammals, and thus reduce raptors from foraging in these habitats, although this may conflict with other road verge wildlife management, and would need careful consideration and further study.

## Acknowledgements

Foremost, I wish to express my sincere appreciation to BTO for providing the data for this project. Additionally, I am extremely thankful to Neil Calbrade, Data Request Coordinator and Bridget Griffin, Ringing & Nest Records Process Manager for communication and assistance with the data request. I am also grateful to the BTO Ringing Scheme and all the ringers that collected the data.

I am deeply grateful to my supervisor Lee Read, MSc International Animal Welfare, Ethics and Law, Programme Leader for Animal Behaviour and Welfare Degree at Kingston Maurward University Centre, for his guidance and continuous support with this project. I also convey my most sincere gratitude to my other supervisor Todd Lewis, PhD, Lecturer in Animal Science at Kingston Maurward University Centre, for his inspiration, dedication, and ongoing assistance with data analysis.

Lastly, I want to thank my family and friends for their endless encouragement.

## Appendices

### Appendix 1. Code Definitions

#### Age Codes

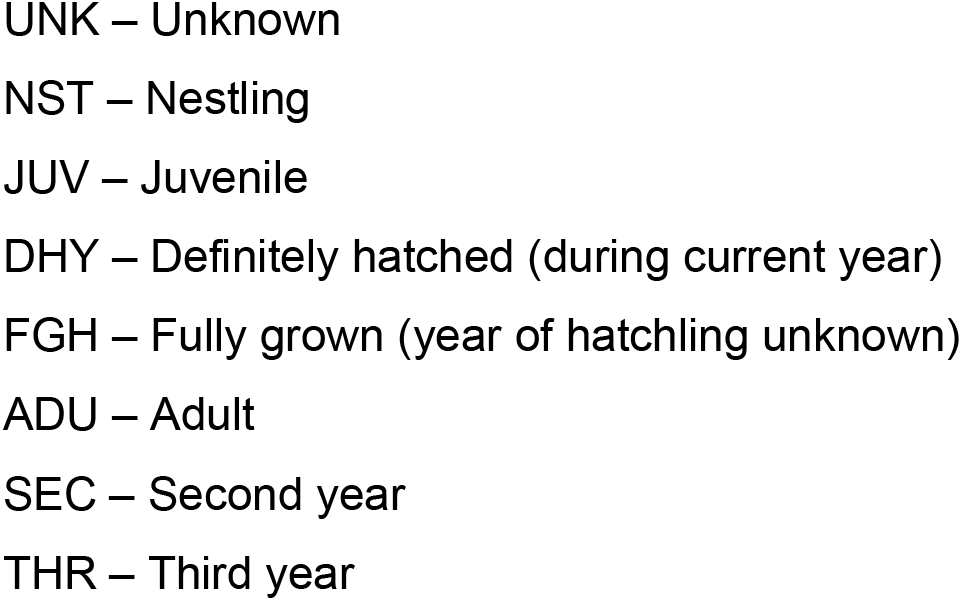

#### Encounter Condition

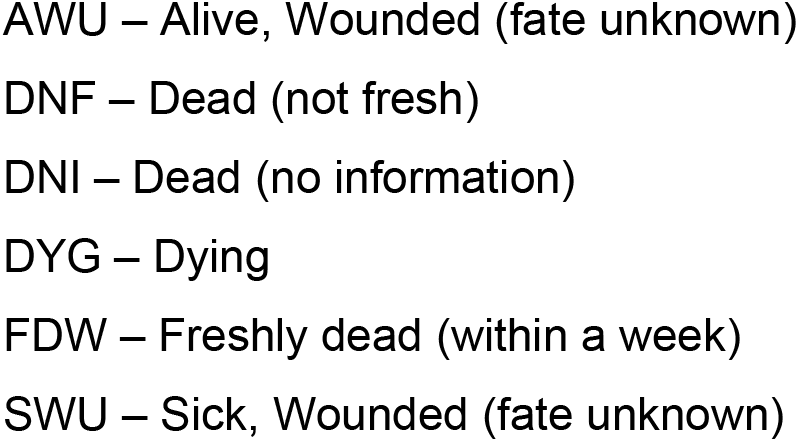

### Appendix 2. JAGS Model Results and Convergence Data

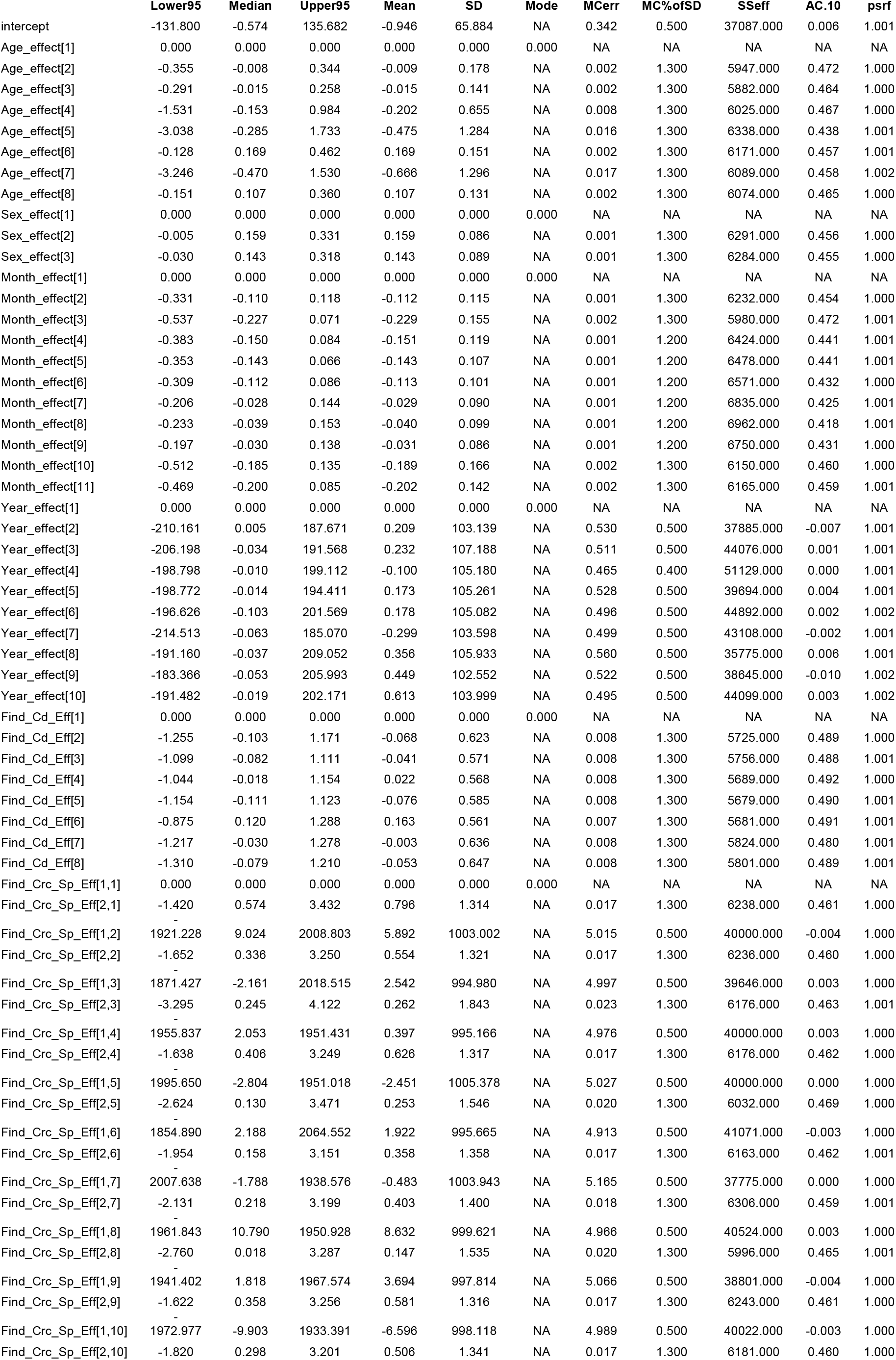
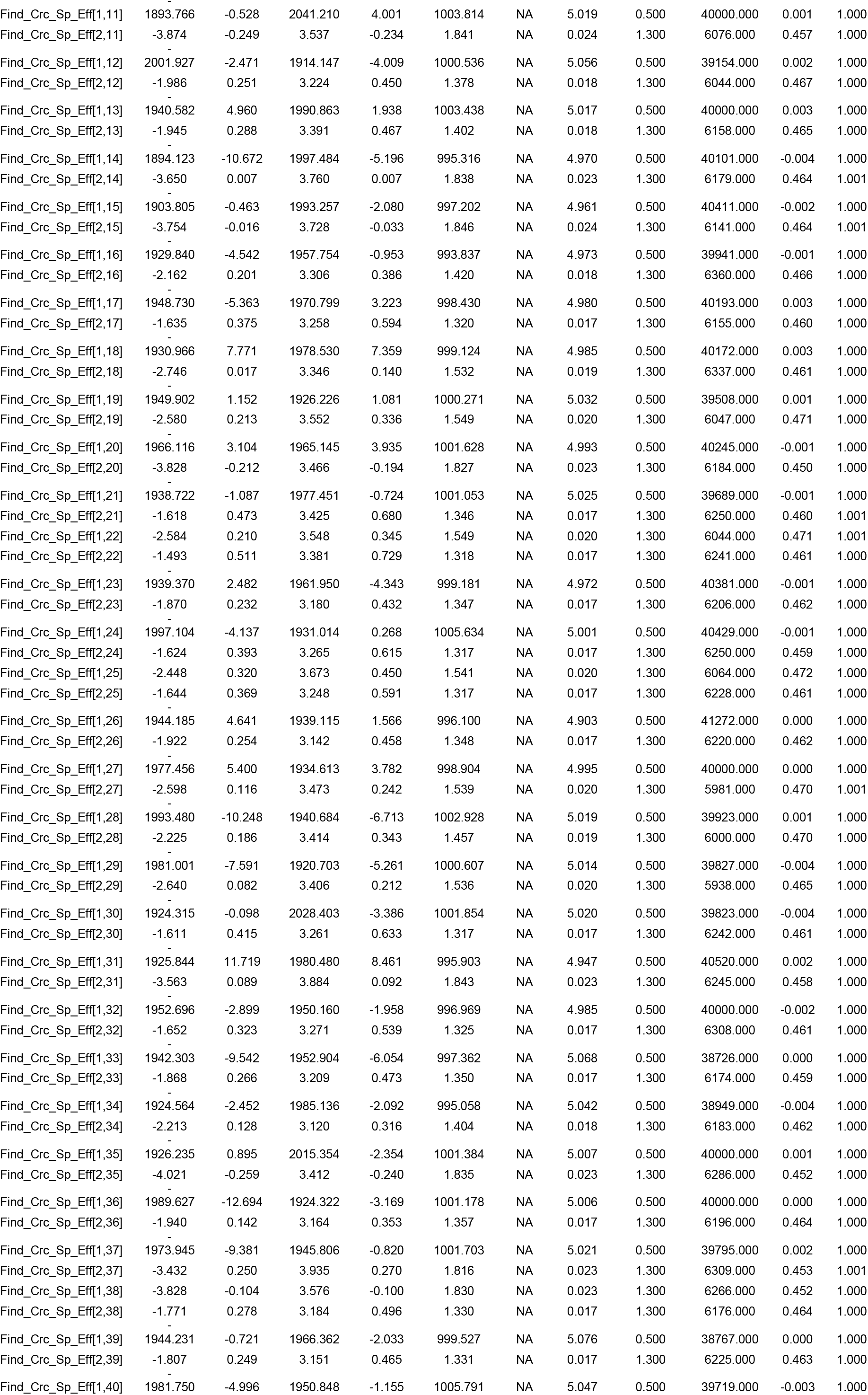
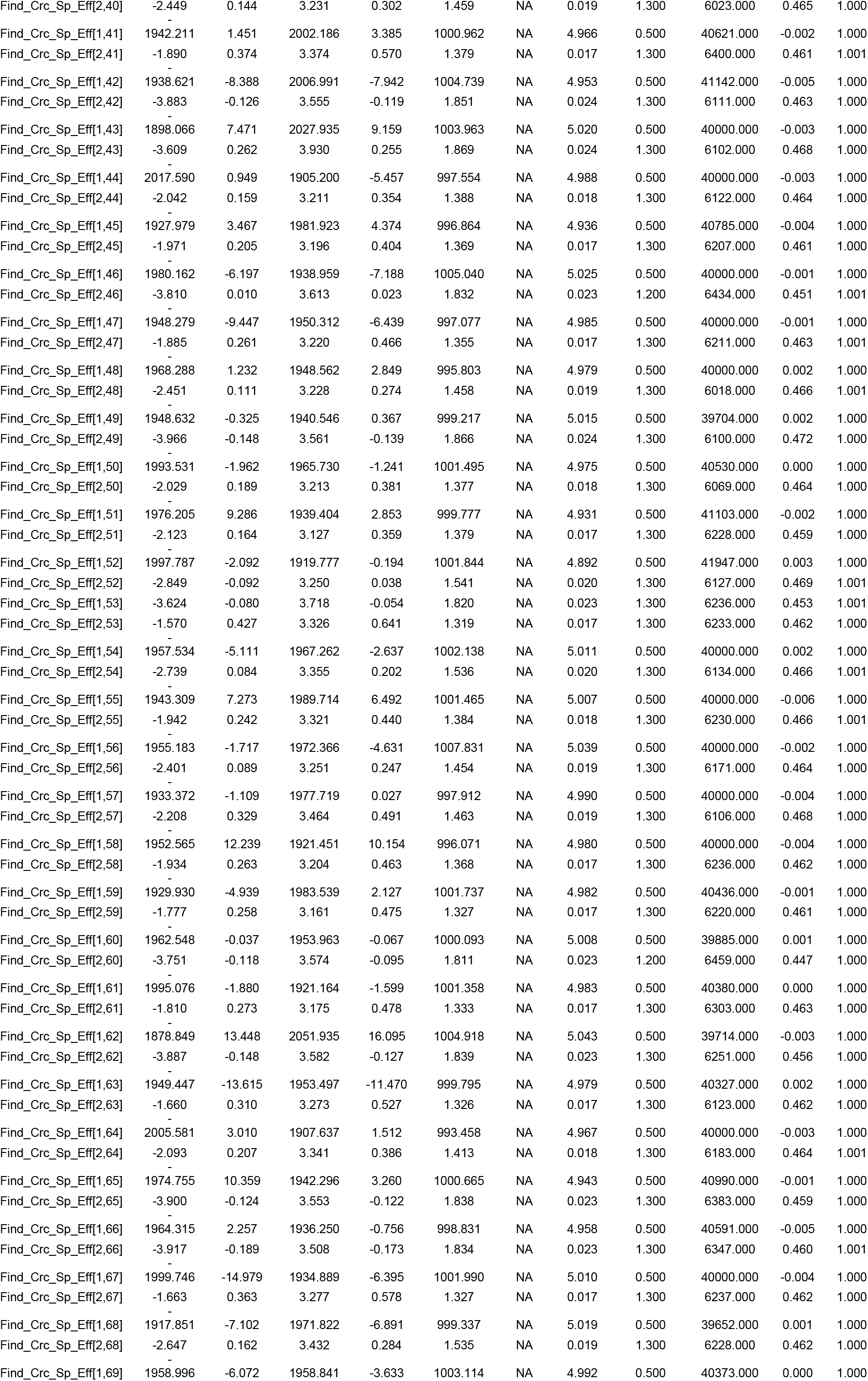
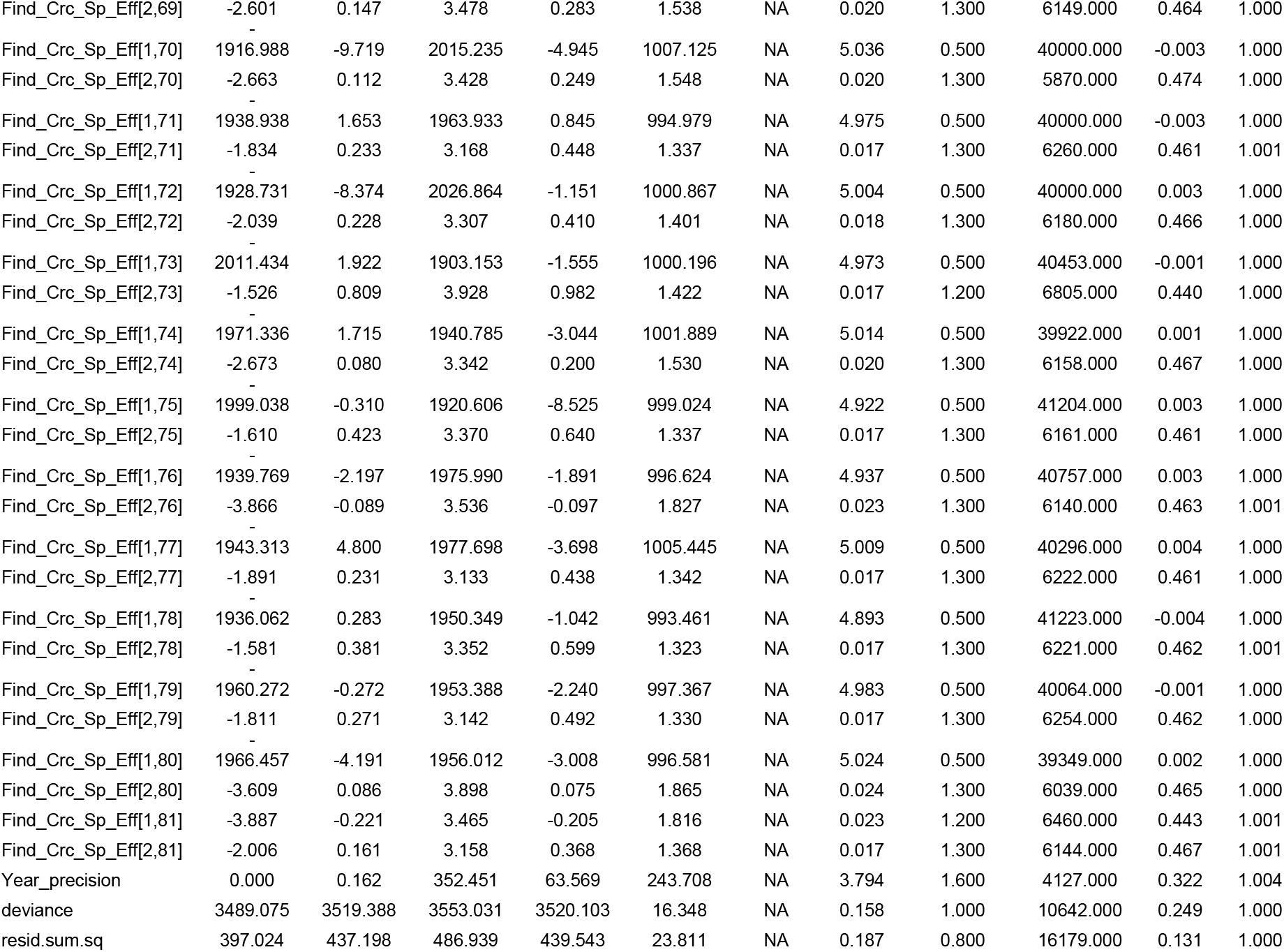

